# An RNA binding polymer specifies nematode sperm fate

**DOI:** 10.1101/088914

**Authors:** Scott Takeo Aoki, Douglas F. Porter, Aman Prasad, Marvin Wickens, Craig A. Bingman, Judith Kimble

**Author notes:** For correspondence: Judith Kimble, HHMI/Department of Biochemistry, University of Wisconsin-Madison, 433 Babcock Drive, Madison, WI 53706-1544. **Abbreviations** RNA: ribonucleic acid DNA: deoxyribonucleic acid FOG: feminization of germline UTR: untranslated region.

## Abstract

Metazoan germ cells develop as sperm or oocytes, depending on chromosomal sex, extrinsic signaling from somatic tissue and intrinsic factors within the germ cells. Gamete fate regulatory networks have been analyzed in nematodes, flies and mammals, but only in *C. elegans* have terminal intrinsic regulators been identified, which include a Tob/BTG protein family member, FOG-3. Canonical Tob/BTG proteins function as monomeric adaptor proteins that link RNA binding proteins to deadenylases. To ask if FOG-3 functions similarly, we first determined its crystal structure. FOG-3 harbors a classical Tob/BTG fold, but unlike other Tob/BTG proteins, FOG-3 dimerizes and these FOG-3 dimers assemble into polymers. The importance of FOG-3 polymers to sperm fate specification was confirmed *in vivo* using CRISPR/Cas9 gene editing to create mutations designed to disrupt the polymer interface. The FOG-3 surface potential is highly basic, suggesting binding to nucleic acid. We find that FOG-3 binds RNA directly with a strong preference for 3’UTRs of oogenic mRNAs. Our results reveal a divergent but striking molecular assembly for proteins with a Tob/BTG fold, make key advances in understanding the mechanism of sperm fate specification and highlight the potential for undiscovered protein polymers in biology.

## Introduction

Metazoan germ cells differentiate as sperm or oocyte. The initial trigger is chromosomal sex, but extrinsic and intrinsic cell fate regulators work subsequently to specify germ cell sex ^reviewed in^ ^1^. Nematode, fly and mammalian germ cells assume a spermatogenic or oogenic fate in response to sex-specific signaling from somatic tissues, and their response relies on intrinsic regulatory factors ^2,3^. Terminal regulators of the sperm/oocyte fate decision have been elusive in most organisms, but have been found in the nematode *C. elegans* ^4,5^. Whereas transcription factors control cell fate in somatic tissues ^e.g. 6,7^, post-transcriptional regulation has emerged as a major mode of control in germ cells. For example, the nematode terminal regulators of germ cell sex determination are conserved post-transcriptional regulators ^8-11^, and RNA regulators also affect germline sex determination in mammals ^12^. A major challenge now is to elucidate the molecular mechanism of these key germ cell fate regulators both *in vitro* and *in vivo.*

An elaborate regulatory network drives the *C. elegans* sperm/oocyte fate decision ^2,13^. Most relevant to this work are the terminal sperm fate regulators FOG-3, a Tob/BTG protein (**Fig. 1a**) ^5,8^, and FOG-1, a cytoplasmic polyadenylation element binding protein (CPEB) ^4,9,10^. Other sperm/oocyte regulators act upstream of these two key regulators. Removal of either the *fog-1* or *fog-3* gene sexually transforms the germline to produce oocytes rather than sperm, the Fog (<u>f</u>eminization of <u>g</u>ermline) phenotype ^4,5^. CPEB and Tob/BTG family members also influence mammalian germ cell development ^14-16^, but a role in the mammalian sperm/oocyte decision is not yet known. Nonetheless, molecular insights into mammalian CPEB and Tob/BTG set the stage for this work. Mammalian CPEB binds specific mRNAs, whereas mammalian Tob/BTG binds CPEB and recruits deadenylases to shorten polyA tails and decrease stability of CPEB target mRNAs ^17,18^. Therefore, these conserved proteins work together to repress mRNA translation ^17^. Similarly, *C. elegans* FOG-1/CPEB and FOG-3/Tob proteins bind each other *in vitro* and co-immunoprecipitate from worms ^11^. In addition, FOG-1/CPEB and FOG-3/Tob associate with a set of common mRNAs and many of their putative targets belong to the oogenesis program, a striking finding since these RNAS were identified in spermatogenic germlines. These results support the model that FOG-1/CPEB and FOG-3/Tob work together in a complex to repress oogenic mRNAs ^11^.

**Figure 1.**
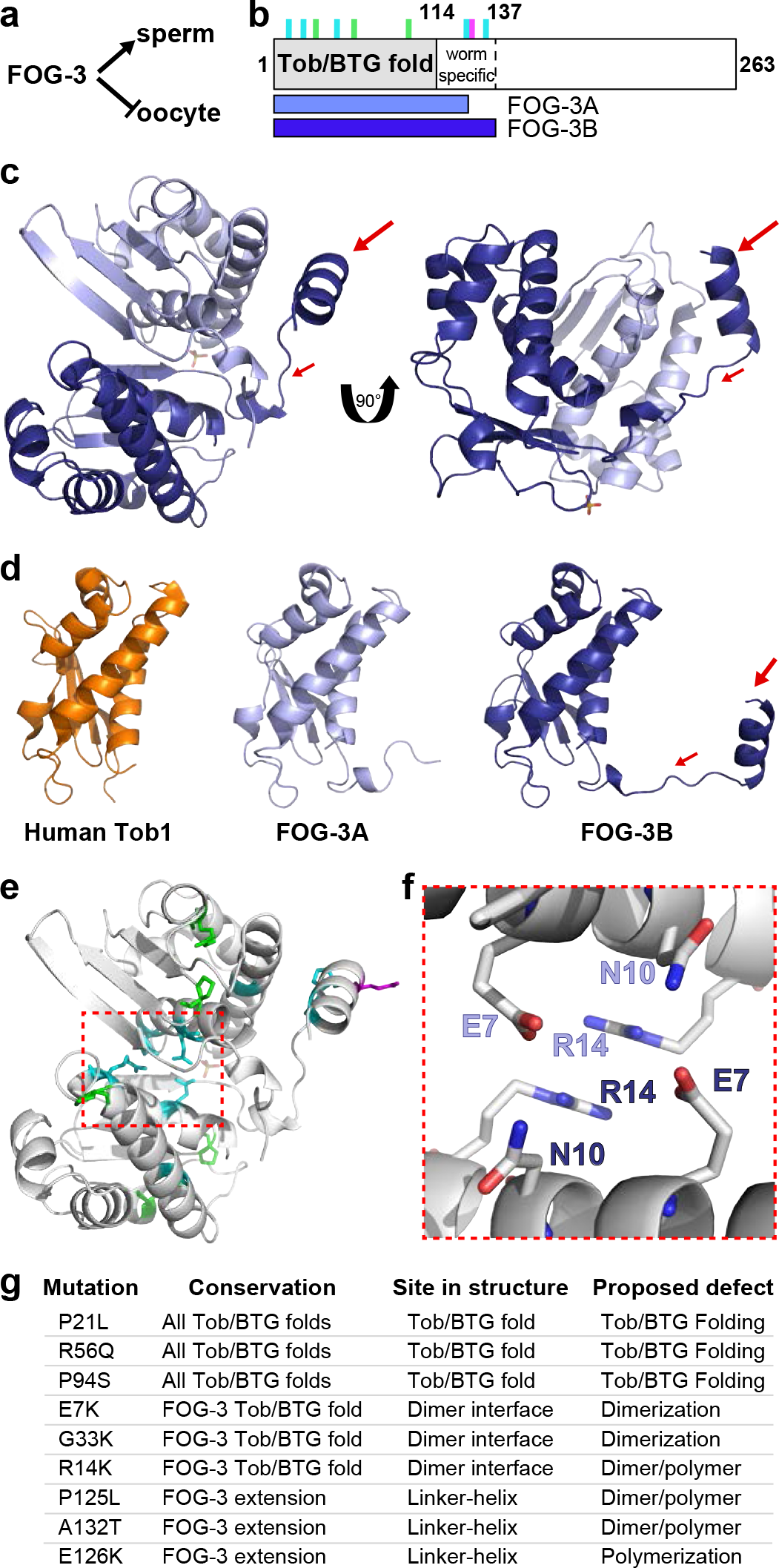
FOG-3 is a divergent Tob/BTG protein. (**a**) FOG-3 specifies the sperm fate. Adult XX and XO animals make oocytes and sperm, respectively. XX animals are essentially females but make sperm transiently as larvae. FOG-3/Tob is a terminal regulator of the sperm fate and essential for sperm fate specification in both XX larvae and XO males. (**b**) Summary diagram of FOG-3. Residues 1-137 are a stable domain (data in **Supplementary Fig. 2**). Predicted Tob/BTG fold is N-terminal (grey); linker-helix forms in the nematode-specific part (worm specific) of the domain (see **Fig. 1c-e**). Above, vertical lines mark sites of missense mutations (also see **Supplementary Fig. 1**): green, residues conserved in all paralogs; cyan, residues conserved only in nematode paralogs; magenta, missense mutation generated in this study. Below, horizontal bars show extents of subunits in crystal dimer, termed FOG-3A (light blue) and FOG-3B (dark blue). (**c**) Asymmetric unit of FOG-3 crystal structure (RCSB PDB ID: 5TD6). FOG-3A (light blue) and FOG-3B (dark blue) co-crystallized with a sulfate (orange). Arrows highlight the nematode-specific linker-helix extension. (**d**) FOG-3 and a human Tob/BTG structures. RMSD of human Tob1 (orange, PDB ID: 2Z15) compared to FOG-3A and B was 1.062 Å and 1.086 Å, respectively. Arrows again highlight the linker-helix extension. (**e**) Location of FOG-3 missense mutants in crystal structure. Color scheme matches that in **b**. Red box outlines region enlarged in **f**. (**f**) Dimer contacts at the Tob/BTG domain interface. Side chains from FOG-3A (light blue) and FOG-3B (dark blue). See text and **Supplementary Fig. 2** for further details. (**g**) Summary of FOG-3 missense mutations.

A battery of *fog-1* and *fog-3* missense mutations, which sexually transform germ cells ^4,5^, provides molecular insights into sperm specification. Most *fog-1* mutations map to the RNA binding domain, which links RNA binding to the FOG-1 role in sperm fate specification ^19^. Similarly, most *fog-3* mutations map to the Tob/BTG fold ^8^ (**Fig. 1b** and **Supplementary Fig. 1**), which links this region to the FOG-3 role in sperm specification. However, few mutations map to residues conserved with mammalian Tob/BTG proteins (**Supplementary Fig. 1**), and no mutations map to residues corresponding to the deadenylase binding interface of human Tob ^20^. We therefore considered the idea that FOG-3 might use a distinct molecular mechanism.

To gain insight into how FOG-3 regulates the sperm fate, we first determined its crystal structure. This structure confirmed the presence of a Tob/BTG fold, but also revealed novel features. Specifically, FOG-3 crystallized as a polymer of dimers. By contrast, canonical Tob/BTG proteins are monomeric ^20,21^. We confirmed FOG-3 polymers with electron microscopy and biochemistry, and demonstrated their biological significance with genetics. The surface potential of the FOG-3 polymer was basic, which suggested RNA binding. Using FOG-3 iCLIP, we confirmed FOG-3 binding and learned that it binds across 3’UTRs of mRNAs belonging to the oogenic program. Together, our results from crystallography, genetics and molecular biology show that FOG-3 functions as an RNA-binding polymer and that FOG-3 targets the 3’UTRs of oogenic transcripts. Our analysis supports a new molecular mechanism for sperm fate specification and highlights the idea that evolution can usurp a well conserved domain to form novel polymers with only modest changes to its primary sequence and structural fold.

## RESULTS

### FOG-3 is a divergent Tob/BTG protein

FOG-3 is predicted to be a Tob/BTG protein. Like canonical members of this family, the FOG-3 primary sequence possesses a predicted N-terminal Tob/BTG domain and a disordered C-terminal region (**Fig. 1b** and **Supplementary Fig. 1**) ^8^. Comparison of FOG-3 sequences from several *Caenorhabditid* species reveals further nematode-specific conservation that extends ~20 amino acids past the predicted Tob/BTG fold. We used recombinant *C. elegans* FOG-3 protein to probe domain boundaries. Full-length recombinant FOG-3 (amino acids 1-263) was unstable, but FOG-3(1-238) was expressed robustly (**Supplementary Fig. 2a,b**). Proteases generated ~15 kDa protected fragments (**Supplementary Fig. 2b**), which by mass spectrometry spanned residues 1-135 after trypsin and 1-142 after elastase cleavage. Both fragments included the predicted Tob/BTG fold plus the nematode-specific extension (**Fig. 1b**). FOG-3(1-137) exhibited a broad elution peak (**Supplementary Fig. 2a**) that could be attributed to a partially unfolded peptide or several multimerization states. Thus, both sequence conservation and *in vitro* analyses suggested that FOG-3 contains a single domain spanning the canonical Tob/BTG fold and a nematode-specific extension.

**Figure 2.**
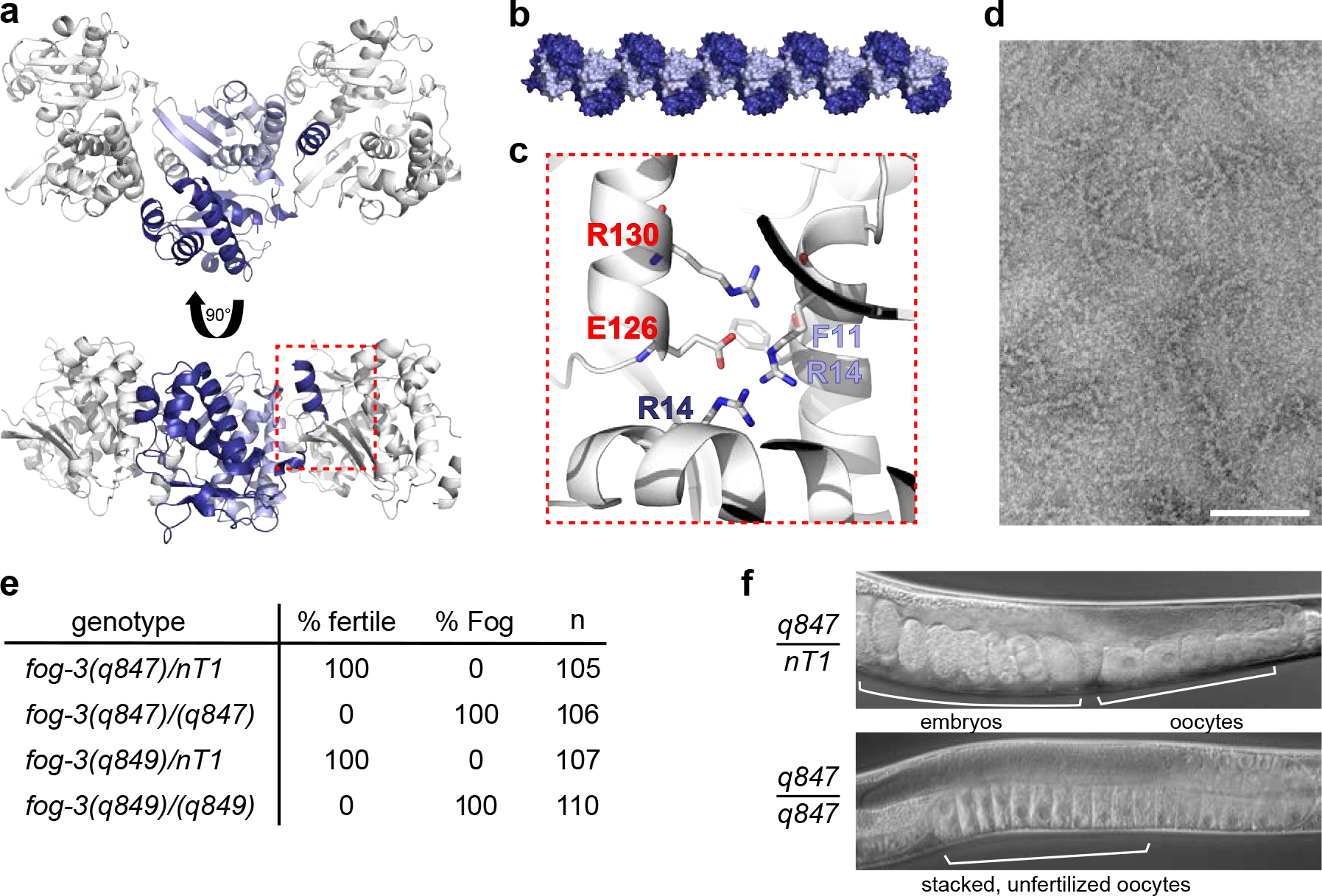
FOG-3 assembles into a polymer. (**a**) Crystal packing of the FOG-3 dimer. FOG-3A and FOG-3B in the asymmetric unit are represented in light and dark blue, respectively. Each dimer buries the helix extension in FOG-3B into the adjacent dimer. The helix extension (red box) is enlarged in **c**. (**b**) Model of FOG-3 polymer, as observed in the crystal. Image generated by extending the crystal symmetry. Subunits colored as in **a**. (**c**) Packing of the helix extension of one FOG-3 subunit into the adjacent dimer. Amino acids in subunits colored as in (**a**), except for red residues from helix extension of the adjacent dimer. (**d**) Negative stain electron microscopy of recombinant FOG-3 1-137. Note the presence of long rods. (**e**) Mutation of a key residue in the helix extension sexually transforms the germline (Fog phenotype). Two identical, but independently generated CRISPR-Cas9 alleles (*q847* and *q849*) mutated glutamate 126 to a lysine (E126K). Alleles were placed over a GFP-expressing balancer (*nT1*). Green (heterozygous) and non-green (homozygous) L4 worms were singled and analyzed 3-4 days later for fertility and the Fog phenotype. (**f**) Representative DIC images of adults heterozygous or homozygous for the E126K mutation. Note the embryos in the heterozygous worms and the oocyte stacking in the homozygous worms.

We pursued the FOG-3 crystal structure to gain insight into this putative Tob/BTG protein. Residues 1-137 gave good yields, but solubility and stability remained issues at high concentration. We improved solubility by retaining the histidine tag and mutating non-conserved residues to amino acids in related FOG-3s (H47N and C117A) (**Supplementary Fig. 1**). Initial crystallization trials proved unfruitful. Thinking that a cofactor was missing, we performed a thermal folding assay ^22^ with various additives (Online Methods). Magnesium and sulfate improved protein thermostability (**Supplementary Fig. 2c**) so we reasoned that they might aid stability during crystallization. FOG-3(1-137, H47N C117A) crystallized in the presence of magnesium sulfate, and these crystals provided a full data set to 2.03 Å (RCSB PDB ID: 5TD6, **Supplementary Table 1**). Phase information was obtained using a human Tob structure (Online Methods).

The FOG-3 crystal structure confirmed the predicted Tob/BTG fold and revealed additional features. Each asymmetric unit contained two FOG-3 subunits, FOG-3A and FOG-3B (**Fig. 1b,c**). For FOG-3A, we could model residues 1-123, missing the last 14 residues, and for FOG-3B we could model nearly the entire peptide chain (1-136) (**Fig. 1b,c**). The structural alignment was excellent between FOG-3 (both A and B) and a previously determined human Tob1 structure (RMSD 1.062-1.084 Å, **Fig. 1d**) ^20^, confirming a Tob/BTG fold in FOG-3. However, unlike other Tob/BTG structures, the FOG-3 structure extended past the classic Tob/BTG fold as a linker-helix extension (**Fig. 1c,d**). Details of the structure supported the idea that the two FOG-3 subunits in the ASU represent a *bona fide* dimer. The buried surface area between the two FOG-3 subunits was large (1034.2 Å) and the interface intricate (**Supplementary Fig. 2d,e**). The linker-helix extension of FOG-3B folded around FOG-3A (**Fig. 1c**), making several hydrogen bonds (**Supplementary Fig. 2d-g**), and the Tob/BTG folds contacted each other at their N-terminal helices, with a hydrogen bond and arginine planar stacking between conserved residues (**Fig. 1e,f**). Dimerization could also explain the elution profile of recombinant FOG-3 (**Supplementary Fig. 2a**). Thus, the crystal structure supported authenticity of the FOG-3 dimer.

### FOG-3 forms a biologically relevant dimer

To ask if the FOG-3 dimer is significant biologically, we analyzed the sites of eight missense mutations that eliminate the ability of FOG-3 to specify sperm fate (**Fig. 1b** and **Supplementary Fig. 1**) ^8^. Our structure included all missense sites (**Fig. 1e,g** and **Supplementary Fig. 1**). Three mutations (P21L, R56Q, P94S) alter residues conserved across Tob/BTG folds, and five others change residues conserved only in FOG-3 and its nematode paralogs (**Fig. 1b,g** and **Supplementary Fig. 1**). The three Tob/BTG fold mutations include two prolines located between helices and an arginine making a hydrogen bond characteristic of Tob/BTG folds (**Supplementary Fig. 2f,h**) ^20,21^. Because of their locations and contacts, we speculate that these residues are crucial to protein folding. Among the other mutations, three (E7K, R14K, G33K) map to the Tob/BTG fold (**Fig. 1f,g**) and two (P125L, A132T) map to the nematode-specific linker-helix extension (**Fig. 1g** and **Supplementary Fig. 2f,i**). Two Tob/BTG fold missense residues (E7K, R14K) belong to the cluster mediating FOG-3 dimerization (**Fig. 1e**) and the third (G33K) maps to a central helix, where a bulky lysine residue could disrupt dimerization via steric hindrance (**Supplementary Fig. 2f,h**). The two mutations outside the Tob fold (P125L, A132T) map to the base and internal face of the link-helix extension in FOG-3B (**Supplementary Fig. 2f,i**), highlighting the importance of this extension to FOG-3 function. We conclude from the structure together with the sites of key missense mutants that FOG-3 likely dimerizes *in vivo* and that dimerization is critical for FOG-3 function.

### Polymerization of FOG-3 dimers

The mapping of two FOG-3 missense mutations to the nematode-specific extension suggested the importance of this linker-helix for sperm fate specification. Such an extension seemed an unusual strategy for dimerization, because the linker-helix interaction is asymmetrical (**Fig. 1c**). We therefore wondered if the linker-helix could be used instead for additional dimer-dimer interactions. To explore this possibility, we extended the crystal symmetry to visualize FOG-3 dimer-dimer interactions in the structure and found the linker-helix extension of each dimer tucked neatly into a cleft in its adjacent FOG-3 dimer (**Fig. 2a**). Each dimer was rotated 180° relative to its neighbor in a continuous pattern to form a polymer within the crystal (**Fig. 2b**). The polymer employs two FOG-3 dimers to complete a 360° turn (**Fig. 2a,b**). Details of the structure suggested that the dimer-dimer interface was authentic. Its surface area was 1112.5 Å (**Supplementary Fig. 2j**), a value similar to that between the dimer subunits, and the interface included 10 hydrogen bonds and four salt bridges (**Supplementary Fig. 2j-l**), with many residues conserved among nematode FOG-3 paralogs (**Supplementary Fig. 1**). For example, the linker-helix of FOG-3B made two salt bridges with the R14 stacking arginines of the adjacent dimer (**Fig. 2c** and **Supplementary Fig. 2l**). The conservation of these interacting residues implies that FOG-3 polymerization is conserved among *Caenorhabditid* species.

We sought to confirm FOG-3 polymerization by complementary *in vitro* methods. The FOG-3 sizing column elution profile gave no hint of polymerization, but polymerization might require a high protein concentration that is not achieved when diluted over the column. Indeed, negative-stain electron microscopy (EM) revealed rods of FOG-3 polymers when recombinant FOG-3(1-137, H47N C117A) was assayed at high concentration (**Fig. 2d**). The appearance of FOG-3 rods was sporadic, making EM an unreliable way to assay polymerization. We turned instead to biochemistry, and used the crystal structure to design lysine mutations at non-conserved residues that could facilitate intra- and inter-dimer crosslinking with the addition of a chemical (**Supplementary Fig. 3a-c**). This FOG-3 “lysine mutant,” which harbored four residues mutated to lysine, could be purified (**Supplementary Fig. 3d,e**) and robustly crosslinked as dimers when incubated with a chemical crosslinker (**Supplementary Fig. 3f**). At higher concentrations of the chemical crosslinker, larger species formed, which we attributed to polymer formation (**Supplementary Fig. 3f**). We next tested crosslinking with that FOG-3 lysine mutant that also harbored a missense mutation (R14K) postulated to impede dimerization. Attempts to crosslink this R14K mutant generated very little dimer and no multimer signal (**Supplementary Fig. 3f**). These results support our structural model that certain FOG-3 missense mutants disrupt dimerization.

**Figure 3.**
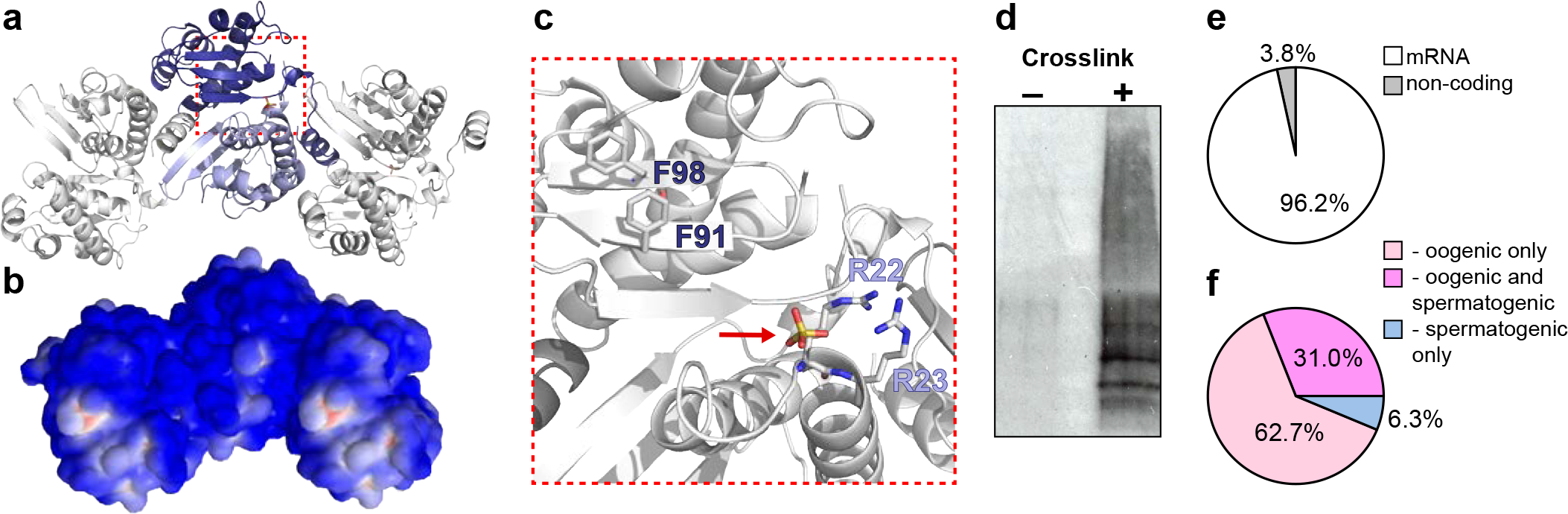
FOG-3 binds directly to mRNAs associated with oogenesis. (**a**) Model of FOG-3 polymer with three dimers; subunits colored as in Figure 1. Region in red box is enlarged in **c**. (**b**) Electrostatic surface potential of polymer modeled in **a**. Blue, basic; red, acidic. (**c**) The bound sulfate (red arrow) is adjacent to the open face of the β-sheet. Amino acids in subunits colored as in **a**. Arginines coordinating sulfate are labeled. Surface of the β-sheet exposes conserved aromatic amino acids, F91 and F98. (**d**) FOG-3 crosslinks with RNA *in vivo*. Worms expressing a rescuing, epitope-tagged FOG-3 transgene were UV-crosslinked (+) or mock treated (-), and FOG-3 immunoprecipitated. Bound sample 5' radiolabeled and run on SDS- PAGE. (**e**) FOG-3 iCLIP enriches for mRNAs. (**f**) Most FOG-3 bound mRNAs belong to the oogenesis program, which includes RNA expressed either only in oogenic germlines (light pink) or in both oogenic and spermatogenic germlines (dark pink), as categorized previously ^11^. See text, Online Methods and **Supplementary Fig. 4** for further details.

No previously identified FOG-3 missense mutants disrupted the dimer-dimer interface specifically. P125L and A132T highlighted the importance of the linker-helix extension for FOG-3 function, but their wild-type residues were not implicated specifically in polymerization. The two R14s of one dimer made contacts with the neighboring dimer, but they were also associated with planar stacking for dimerization (**Fig. 1e**). Thus, we lacked a mutation that disrupts polymerization specifically. The FOG-3 structure revealed a glutamate (E126) on the solvent exposed surface of the linker-helix that contacts the neighboring dimer (**Fig. 2c**). We reasoned that changing E126 to a positively charged amino acid (E126K) would disrupt polymerization. Indeed, chemical crosslinking of a FOG-3 lysine mutant with an E126K mutation had little effect on dimer crosslinking, but prevented formation of higher ordered multimer species (**Supplementary Fig. 3f**). We conclude that FOG-3 dimer-dimer contacts facilitate polymerization. To test the importance of polymerization to sperm fate specification *in vivo*, we used CRISPR-Cas9 gene editing ^23, see Online Methods^ to introduce the E126K change in the endogenous *fog-3* gene. Animals homozygous for either of two independent E126K alleles were unable to make sperm and instead had a fully penetrant Fog phenotype (**Fig. 2e,f**). Therefore, FOG-3 is likely to function as a polymer to promote sperm fate.

### FOG-3 binds RNA directly across 3’UTRs of oogenic-associated transcripts

Our crystal structure challenged the idea that nematode FOG-3 regulates its target RNAs via the mechanism elucidated for mammalian Tob/BTG proteins. Canonical Tob/BTGs function as monomeric adapters to recruit deadenylases to RNA binding proteins and their target mRNAs ^24^. However, several structural features suggested that FOG-3 might bind RNA directly rather than serving as an adapter between proteins. The electrostatic surface potential of the FOG-3 polymer is highly basic (**Fig. 3a,b**) and conserved aromatic residues (F91 and F98) are accessible at the surface with potential for base-stacking to nucleic acid (**Fig. 3c** and **Supplementary Fig. 1**). Moreover, a sulfate co-crystallized with FOG-3 between adjacent polymers (**Fig. 3a,c**), which appeared to balance the positive charge of conserved cationic residues R22 and R23 (**Fig. 3c** and **Supplementary Fig. 1**). We speculate that this sulfate may mark where FOG-3 binds the phosphate backbone of nucleic acid.

We tested whether FOG-3 binds directly to RNA in sperm-fated nematode germ cells. FOG-3 immunoprecipitation (IP) enriched for radiolabelled RNA after UV crosslinking (**Fig. 3d**, **Supplementary Fig. 4a-c**), a treatment that creates covalent bonds between proteins and RNA ^25^. The strong radiolabelled signal versus control provided evidence that FOG-3 binds RNA directly. We then used iCLIP 26 to identify RNAs crosslinked to FOG-3 (**Supplementary Fig. 4c-f**) and to learn the location of where it binds within those RNAs. After normalization to a negative control (see Online Methods), FOG-3 enriched for 955 mRNA targets and 38 non-coding RNAs (**Fig. 3e**, **Supplementary Table 2** and **Supplementary Table 3**). Remarkably, ~94% of the mRNA targets belonged to the oogenesis program (**Fig. 3f** and **Supplementary Fig. 6g**), despite their immunoprecipitation from sperm-fated nematode germ cells. The RNA targets identified by iCLIP overlapped with an earlier list of FOG-3 targets, identified by microarray (**Supplementary Fig. 6g,h**, p value < 10^-208^), but iCLIP significantly increased enrichment for oogenic RNAs and decreased spermatogenic RNAs (**Supplementary Fig. 6g,h**). Because iCLIP is more stringent than probe-based microarray methods for identifying targets ^27^, we suggest that iCLIP improved the signal-to-noise ratio of RNAs that immunoprecipitated with FOG-3 from these sperm-fated germ cells. We conclude that FOG-3 binds directly to targets that belong largely to the oogenesis program.

We mapped the sites of FOG-3 binding within its targets. A majority of its binding sites mapped to 3’UTRs (**Fig. 4a,b** and **Supplementary Fig. 4i,j**). Few peaks were observed in the 5'UTR and coding regions (**Fig. 4a,b**), implying that FOG-3 binding is largely restricted to 3’UTRs. If FOG-3 binds RNA as a polymer, it should leave an extensive footprint. Consistent with this idea, 624 of 955 protein-coding genes (65.3%) had two or more sequence peaks that spanned 3’UTRs (**Fig. 4c,d** and **Supplementary Fig. 4i-k**). This binding pattern is reminiscent of 3’UTR multi-site RNA binding proteins, like HuR ^28^. Gaps between peaks might signify authentic absences of FOG-3 binding or they might represent preferred sites of enzymatic digestion during iCLIP (see Methods). We sought motifs enriched near FOG-3 binding sites and found enrichment of a CUCAC motif (**Supplementary Fig. 4l,** p value < 1.8 x 10^-229^). CUCA is part of the GLD-1/STAR signature motif ^29^. GLD-1 regulates germline sex determination, but it can promote either the sperm or oocyte fate ^30^. We conclude that FOG-3 binds across 3’UTRs of its target mRNAs and suggest that it specifies the sperm fate by repressing mRNAs in the oogenic program.

**Figure 4.**
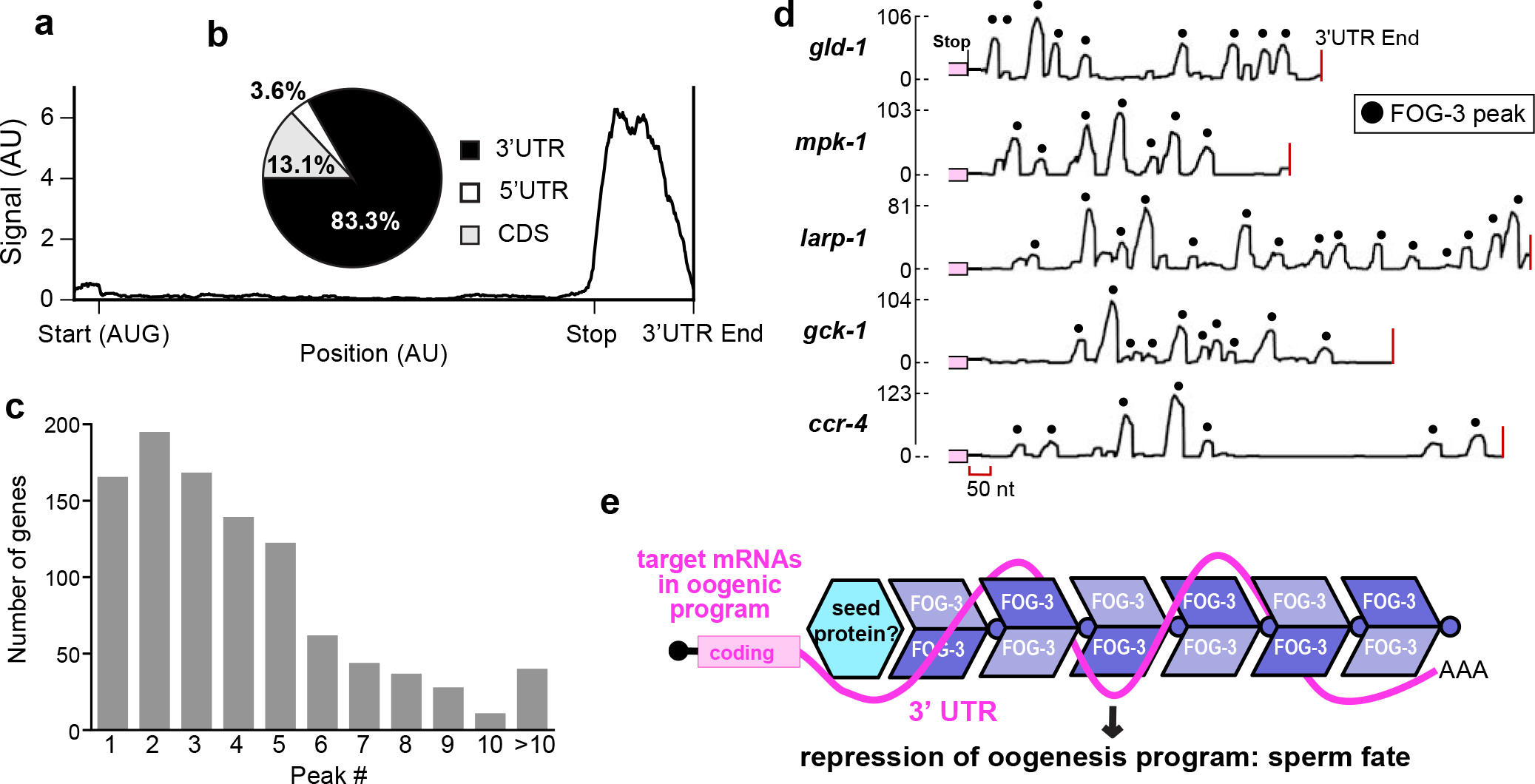
FOG-3 binds 3’UTRs *in vivo*. (**a**) Distribution of FOG-3 iCLIP sequence reads within mRNAs. Transcript lengths normalized for 5'UTRs, Coding Sequences (CDS) and 3’UTRs (50, 1000 and 200 nucleotides, respectively) are reported as arbitrary units (AU). (**b**) Percentages of reads in mRNA regions. (**c**) Many FOG-3 targets possess multiple binding peaks, defined as described (see Online Methods). For top mRNA targets, binding sites span their 3’UTR, as shown in **d**. (**d**) Examples of FOG-3 binding peaks across 3’UTRs. X-axis, 3’UTR with coding region in pink and 3’end marked by red line; y-axis, number of mapped reads. Peaks are marked by black dots; their heights correspond to number of mapped reads. (**e**) Model. FOG-3 polymer binds across 3’UTRs and promotes the sperm fate by repressing mRNAs in the oogenic program. FOG-3 may find its specific target mRNAs by interacting with another RNA binding protein. Further discussion in the main text.

## DISCUSSION

Regulation of gene expression lies at the heart of cell fate specification. Here we connect a poorly understood cell fate decision, specification of the sperm fate, with an unexpected RNA-binding regulatory polymer. We find that the FOG-3 protein can assemble as a polymer *in vitro*, that its polymerization is critical for sperm fate specification *in vivo* and that FOG-3 binds a battery of oogenic mRNAs directly, preferentially at multiple sites throughout their 3’UTRs. These advances into understanding the molecular basis of the sperm fate decision were made possible by solving the crystal structure of a key regulatory protein that then could be queried with missense mutations and CRISPR/Cas9 gene editing to test the importance of structural features. Our findings support a model in which the FOG-3 polymer binds across 3’UTRs to regulate the sperm fate (**Fig. 4e**). Our model further suggests that mRNAs may wrap around the FOG-3 polymer, an idea based on speculation that the positions of co-crystallized sulfates may be potential mRNA phosphate backbone binding sites. By this reasoning, the FOG-3 polymer would package its target mRNAs (**Fig. 4e**). This model is reminiscent of viral assembly proteins that form helical polymers to package RNA genomes ^31^. Another example is Bicaudal-C (bicc1), which multimerizes via its SAM domain to regulate RNA localization and translational silencing ^32^. In our study, recombinant FOG-3 alone was not sufficient to robustly form polymers at low protein concentrations (**Supplementary Fig. 2a**). Like viral assembly proteins and other nucleic acid binding polymers ^33^, FOG-3 may require an mRNA scaffold to form collaborative filaments.

Our analysis of FOG-3 mRNA targets greatly improves our understanding of how FOG-3 regulates sperm fate specification. First, a previous study seeking FOG-3 associated mRNAs used an older technology with microarrays 11 and did not find as convincing an enrichment for oogenic mRNAs (**Supplementary Fig. 6g,h**). Our use of iCLIP demonstrates that virtually all (~94%) FOG-3 targets belong to the oogenic program (**Fig. 3f**), even though spermatogenic germ cells were used to find the FOG-3 targets. Second, it was not known prior to this work that FOG-3 is itself an RNA-binding protein and therefore it could not be known where it binds. Here we demonstrate that FOG-3 not only binds directly to RNA but it binds to 3’UTRs of its target mRNAs. A common 3’UTR regulatory mechanism involves recruitment of RNA-modifying enzymes to regulate mRNA expression ^34,35^. Indeed, mammalian BTG/Tob proteins regulate their targets in this way ^17,18^. Our discovery of FOG-3 polymerization suggests a different mechanism. FOG-3 lacks conservation at sites corresponding to the mammalian Tob/deadenylase binding interface ^20^, arguing against FOG-3 recruitment of a deadenylase. We suggest that instead FOG-3 binds as a polymer across 3’UTRs to either preclude other 3’UTR protein binding or to localize its targets for repression. We do not yet know if FOG-3 provides RNA binding specificity or relies on another RNA-binding protein to seed its polymerization (**Fig. 4e**). If the seed is critical, the most likely candidate is FOG-1/CPEB, which binds FOG-3 and associates with a set of common oogenic mRNAs ^11^. Another candidate might be the GLD-1 STAR/Quaking RNA binding protein, an idea based on the enrichment in FOG-3 target 3’UTRs for the core GLD-1 RNA binding motif ^29^ (**Supplementary Fig. 6i**). Because GLD-1 influences both sperm ^36^ and oocyte specification ^30^, it might seed FOG-3 polymers to drive the sperm fate or compete with FOG-3 binding to drive the oocyte fate. A third possibility is that FOG-3 self-assembles on its RNA targets via either sequence or RNA secondary structure motifs that we could not detect.

Polymerization appears to be a unique feature of FOG-3-related Tob/BTG proteins. The critical residues at the dimer interface are not conserved in mammalian Tob/BTG proteins (**Supplementary Fig. 1**) and these mammalian homologs show no evidence of dimerization or polymerization ^20,21^. Because only a few Tob/BTG proteins have been biochemically characterized, FOG-3 might yet have a mammalian counterpart. However, given the sequence differences between nematode and mammalian Tob/BTG proteins, we favor instead the idea that an existing protein fold was adapted during evolution to generate a polymer. That evolution from monomer to polymer required the generation of inter-subunit and inter-dimer interacting surfaces. In the case of FOG-3, the inter-dimer interface was created by adding a linker-helix extension that fits into the cleft of its neighbor, adding a binding surface to one side of the dimer. This extension provides directionality for polymer assembly. Intriguingly, other nucleic acid binding polymers use a similar strategy. The RNA-binding proteins that package the genomes of RNA viruses possess a core folded domain plus a C-terminal linker-helix or linker-β-sheet extension to drive multimerization ^31^. Similarly, adenovirus E4-ORF3 oncoprotein forms a polymer with its central core dimer and C-terminal linker-β sheet extension ^37^. Our discovery of the FOG-3 polymer was surprising given models for mammalian Tob/BTG proteins, and provides an example of how protein domains can form covert assemblies for novel roles in RNA regulation and cell fate.

## METHODS

Methods and any associated references are available in the online version of the paper.

## ACKNOWLEDGMENTS

The authors thank R. Massey for electron microscopy image acquisition; M. Cox for equipment; A. Helsley-Marchbanks for help preparing the manuscript; L. Vanderploeg for help with the figures; and members of the Kimble and Wickens labs for helpful discussions. Use of the Advanced Photon Source was supported by the U.S. DOE under Contract No. DE-AC02-06CH11357. Use of the LS-CAT Sector 21 was supported by the Michigan Economic Development Corporation and the Michigan Technology Tri-Corridor (Grant 085P1000817). GM/CA@APS has been funded in whole or in part with Federal funds from the National Cancer Institute (ACB-12002) and the National Institute of General Medical Sciences (AGM-12006). STA was supported by NIH grants F32HD071692 and K99HD081208. CAB was supported by NIH grants GM094584, GM094622 and GM098248. MW was supported by NIH grant GM50942. JK is an Investigator of the Howard Hughes Medical Institute.

## AUTHOR CONTRIBUTIONS

STA designed and performed experiments, analyzed the data and wrote the paper; DFP and AP designed and performed experiments and analyzed the data; CAB acquired and analyzed data; MW analyzed the data and drafted the manuscript; JK analyzed the data and wrote the paper.

---

**Supplementary Figure 1:**
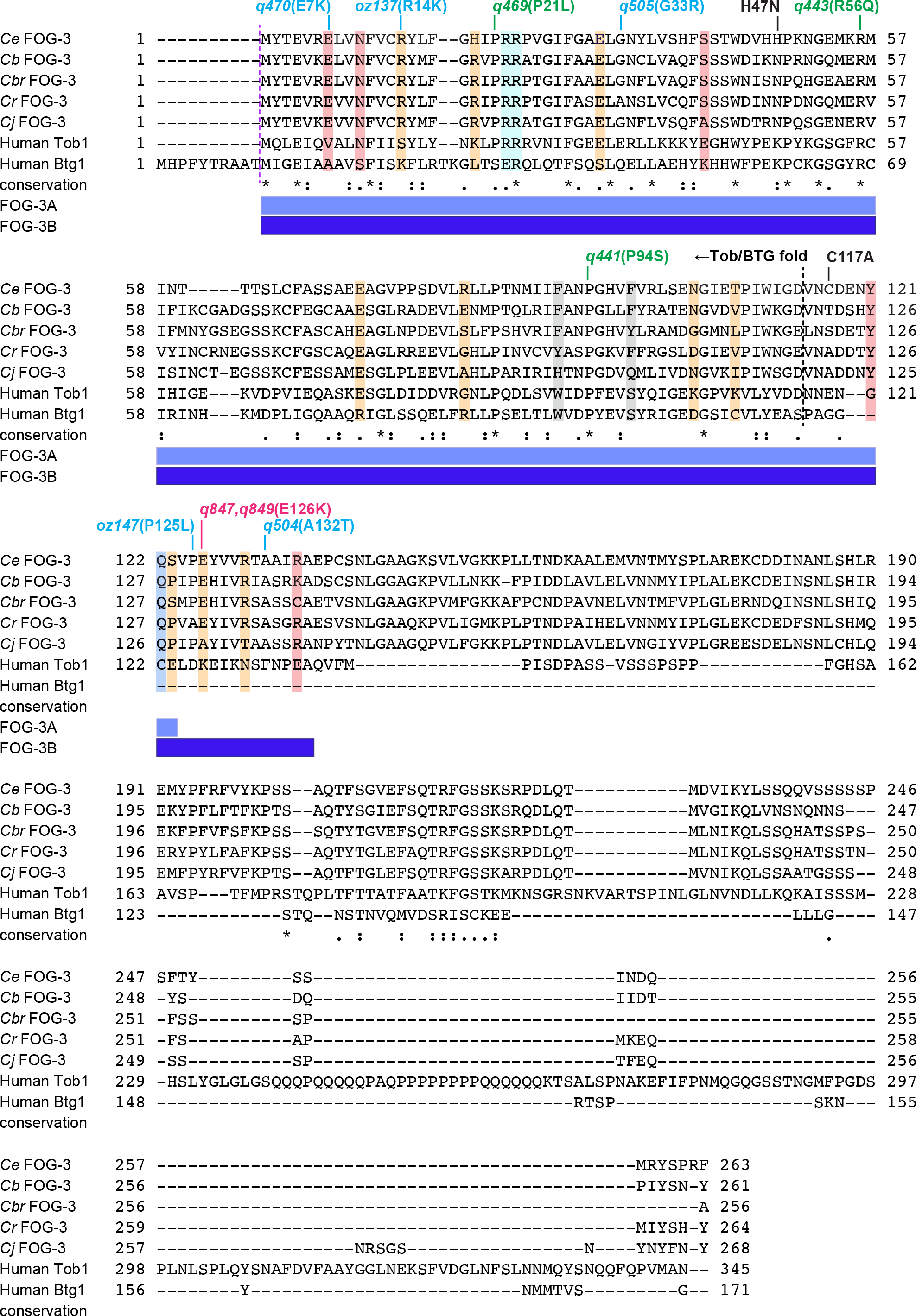
Alignment of FOG-3 ortholog sequences. Amino acid sequence alignment of FOG-3 orthologs, including human Tob and BTG proteins. Nematode orthologs include FOG-3 from *C. elegans* (*Ce*), *C. briggsae* (*Cb*), *C. brenneri* (*Cbr*), *C. remanei* (*Cr*), and *C. japonica* (*Cj*). Alignment by T-coffee ^38^. Conservation noted by identity (*) plus high (:) or moderate (.) similarity. Missense alleles are labeled with their amino acid changes ^8; this work^; allele colors mark conservation among most orthologs (green), conservation among most nematode orthologs (blue) and a mutation generated in this study (magenta). Boundary of the canonical Tob/BTG fold is marked with a dashed line; extents of dimer subunits are shown below, including subunit A (light blue) and subunit B (dark blue). Amino acids highlighted in red, orange and blue indicate inter-dimer contacts, dimer-dimer contacts, and both, respectively; these contacts include both hydrogen bonds and salt bridges. Amino acids in grey are found at the solvent accessible face of the beta sheet and those in cyan are the side chains coordinating the sulfate.

**Supplementary Figure 2:**
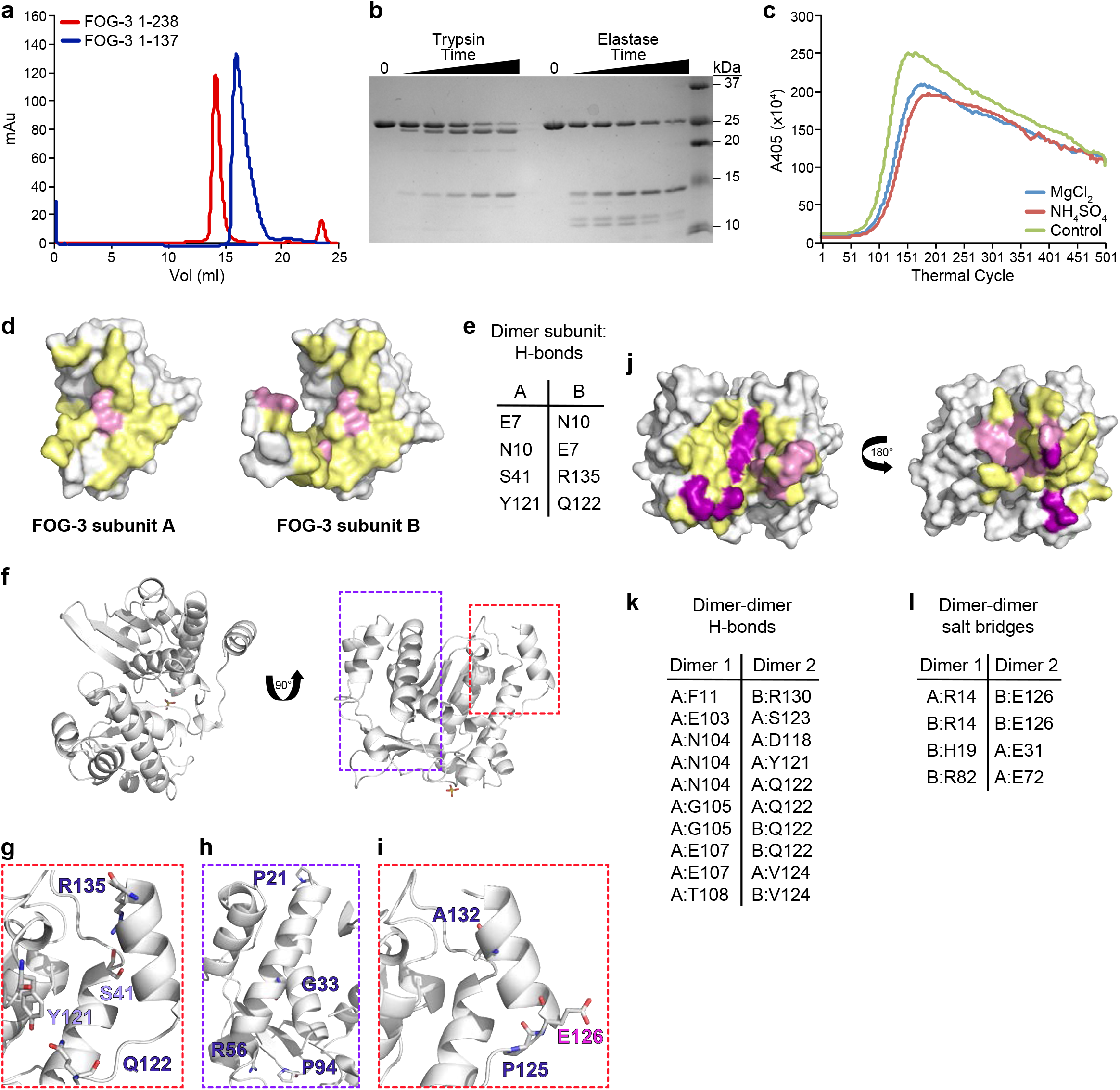
FOG-3 biochemical characterization, crystal structure protein contacts and missense mutants. (**a**) Size exclusion chromatography elution profile of recombinant FOG-3 protein. Red, amino acids 1-238 with its histidine tag; blue, amino acids 1-137 with histidine tag. A280 milliabsorbance units, mAU. (**b**) Mapping a FOG-3 Tob/BTG-containing domain. FOG-3 1-238 was incubated with either trypsin or elastase, and samples were collected over time. Incubation with either protease produced a cleavage product of ~15 kDa. Protein incubated without protease labeled as “0.” (**c**) Thermal folding assay reveals domain stabilization with magnesium and sulfate. See methods and text for further details. (**d**) Interacting residues between FOG-3 subunits. Pink, residues making hydrogen bonds; yellow, other contacting residues, based on distance. (**e**) Table of subunit-subunit hydrogen bonds. (**f**) FOG-3 dimer with boxed regions enlarged in **g-i**. (**g-i**) Residues from FOG-3A are colored light blue and those from FOG-3B are dark blue. (**g**) Inter-subunit contacts made with linker-helix extension in FOG-3B. (**h**) Sites of *fog-3* missense mutants conserved across most FOG-3 orthologs, including human Tob and BTG proteins. (**i**) Sites of *fog-3* missense mutants in linker-helix extension, including one generated in this study (magenta). (**j**) Interacting residues between FOG-3 dimers reveal a dimer-dimer footprint. Salt bridges, hydrogen bonds, and contacting residues, deduced from proximity, are labeled in magenta, pink, and yellow, respectively. (**k**) Table of dimer-dimer hydrogen bonds. (**l**) Table of dimer-dimer salt bridges.

**Supplementary Figure 3:**
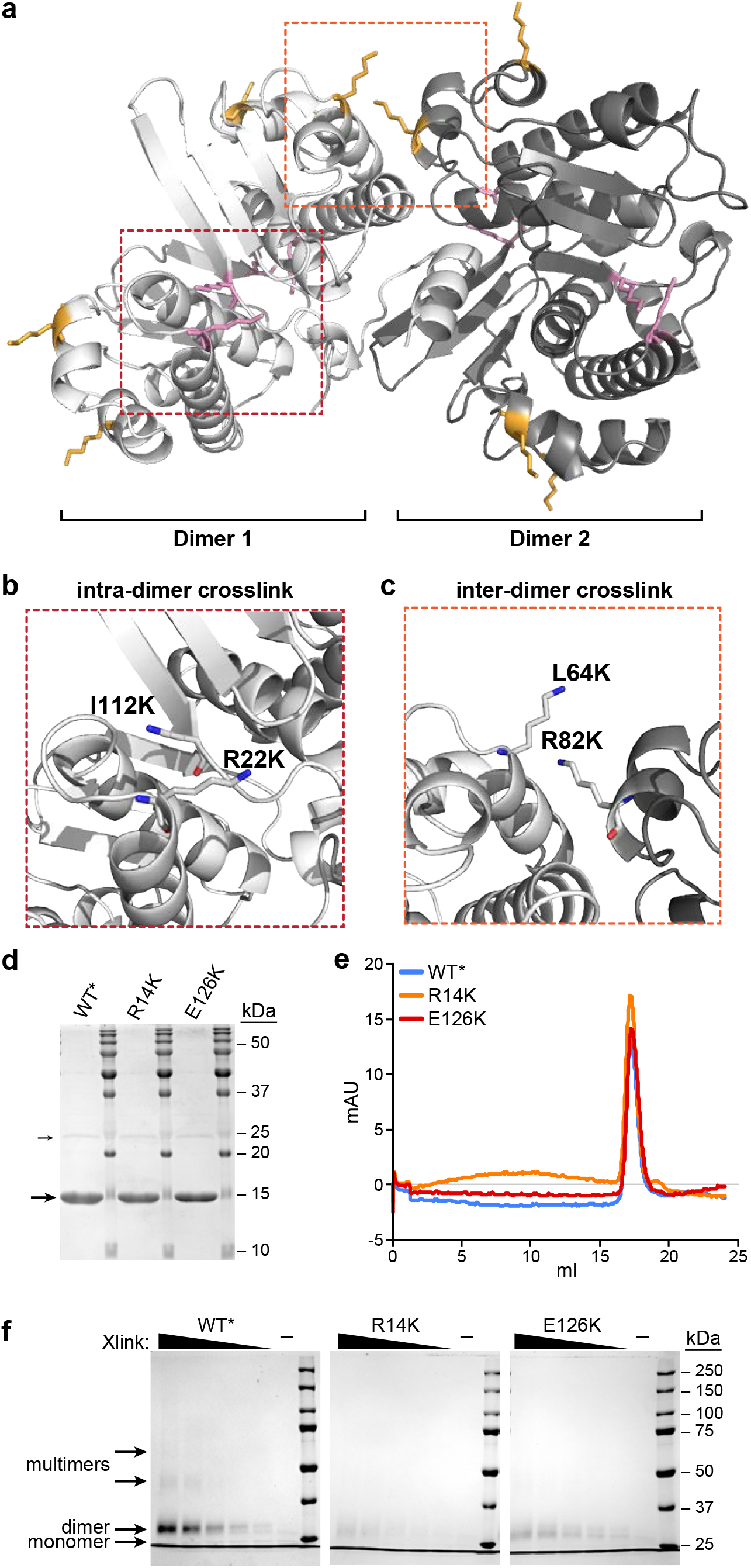
Chemical crosslinking traps FOG-3 polymers *in vitro*. (**a-c**) Lysine mutagenesis of recombinant FOG-3 generates sites permitting intra- and inter-dimer crosslinks. Location of lysine mutations in the context of two FOG-3 dimers (**a**), which are enlarged in **b** and **c**. Red box, lysine mutations R22K and I112K (pink) facilitate an intra-dimer crosslink. Orange box, lysine mutations L64K and R82K (gold) facilitate an inter-dimer crosslink. (**d**) Coomassie stained gel of purified FOG-3 recombinant proteins. Left, “wild-type” (WT*) FOG-3 (1-140, H47N C117A) with lysine mutations; middle, missense mutant R14K predicted to abrogate dimer formation; right, missense mutant E126K predicted to abrogate polymer formation. All proteins (10 μg each) ran at ~15 kDa (large arrow). In addition, a minor ~25 kDa contaminant was observed (small arrow), which was also seen after crosslinking (see **f**). (**e**) Size exclusion chromatography elution profile of WT*, R14K, and E126K recombinant FOG-3 (80 μg). A280 milli-absorbance units, mAU. (**f**) Coomassie stained gel of recombinant protein incubated with increasing amounts of BS3 crosslinker. "-" represents no BS3 included.

**Supplementary Figure 4:**
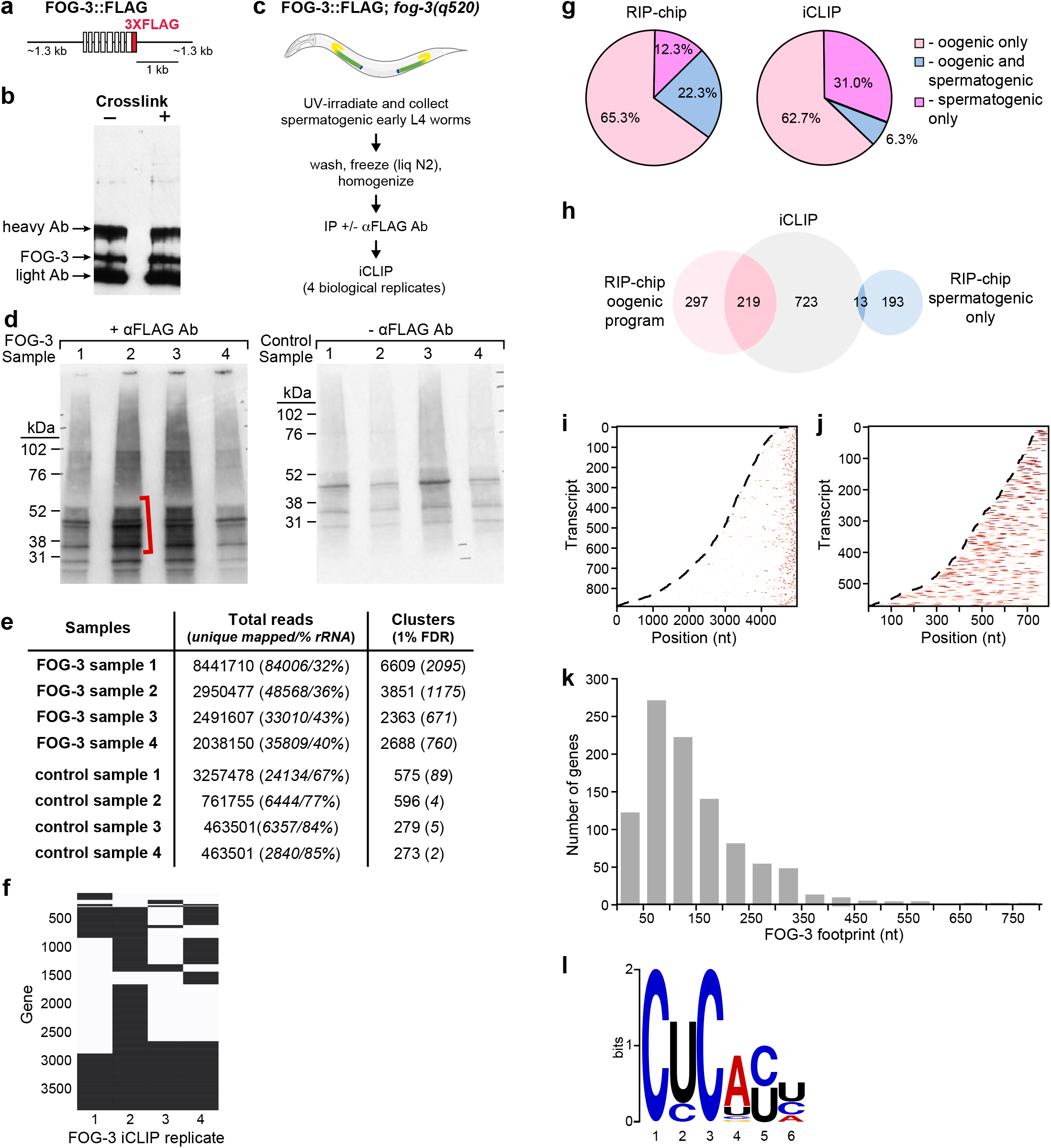
iCLIP supplement. (**a**) Diagram of FOG-3::FLAG transgene, adapted from ^11^. (**b**) Immunoblot of FOG-3 immunoprecipitation samples, visualized with αFLAG antibody. (**c**) Diagram of FOG-3 iCLIP protocol, adapted from ^11^. (**d**) Gel analysis of samples used for iCLIP. Each sample was immunoprecipitated with (+) or without (-) αFLAG antibody, radiolabeled and run on the SDS-PAGE gel. The region above the expected size for FOG-3::FLAG (red bracket) was used for iCLIP processing and sequencing. Samples include four biological replicates and their paired controls. (**e**) FOG-3 iCLIP reads and cluster (regions of overlapping reads) statistics. Unique mapped reads (*middle column*) were determined by mapping with STAR, filtering out multimapping reads and low confidence alignments, and collapsing duplicate reads (see Online Methods). The fraction of unique mapped reads that mapped to rRNA is given as a percentage. The number of significant clusters at FDR 1% (right column) is highly dependent on FOG-3 purification. See Online Methods for further details. (**f**) Targets (black) identified for separate FOG-3 replicates. (**g**) Comparison of FOG-3 targets identified using FOG-3 iCLIP versus FOG-3 RIP-chip (microarray). Targets belonging to the oogenic program include mRNAs found only in oogenic germlines as well as mRNAs found in both oogenic and spermatogenic germlines, as described ^11^. (**h**) Venn diagram of mRNA target overlap between FOG-3 iCLIP (this study) and FOG-3 RIP-chip (FOG-3 IP with microarray analysis of associated RNAs) ^11^. (**i,j**) FOG-3 binding sites are represented on a heat map, from no signal (white) to strong signal (red). Only genes with annotated 3’UTRs of at least 50 nt were included. (**i**) FOG-3 binding sites are at the 3’end of mRNAs, which are arranged by predicted nucleotide (nt) length from 5’ to 3’. Dashed line marks the 5’ end. Note prevalence of FOG-3 binding sites at the 3’ ends of transcripts. (**j**) Binding sites occur throughout 3’UTRs, which are arranged by predicted 3’UTR nucleotide (nt) length from stop codon (dashed line) to 3’ end. (**k**) FOG-3 iCLIP footprint on target transcripts. Coverage includes transcript regions above two reads deep. (**l**) Motif analysis of iCLIP clusters. Analysis and image generated by MEME ^39^, except T was replaced by U.

## Materials and Methods

### Biochemistry and Crystallography

#### Protein expression and purification

Full length FOG-3 (1-263) was amplified from *C. elegans* N2 cDNA with primers that included a six-histidine tag, stop codon and 12 nucleotides suitable for annealing with ligation independent cloning (LIC^1^). Mixed stage N2 cDNA was generated by reverse transcription (SuperScript® II Reverse Transcriptase, Thermo Fisher) with oligo-dT (Ambion). The FOG-3 PCR product was cloned into a pET21a (EMD Millipore) bacterial expression plasmid by LIC. From this plasmid, histidine-tagged FOG-3 (1-238) and FOG-3 (1- 137 H47N, C117A) was amplified by PCR and inserted into pET21a by Gibson cloning^2^. Expression plasmids were transformed into Rosetta™2(DE3) cells (EMD Millipore), grown in LB (MP Biomedicals) for 5 hours at 37°C until A600 = ~0.8. The culture was then induced with 0.1 mM IPTG (MP Biomedicals), and grown at 16°C for 16-20 hours prior to collection, centrifugation, washing and freezing in liquid nitrogen. Cells were defrosted on ice and reconstituted in lysis buffer (20 mM NaPO_4_ pH 7.4, 300 mM NaCl, 10 mM imidazole, 5 mM β-mercaptoethanol) with cOmplete protease inhibitors (Roche). Cells were lysed with a French Press, centrifuged (3220 *x g* and 10000 *x g*) to remove unlysed cells and precipitate, and incubated with Nickel-NTA beads (Thermo Scientific) for 2 hours at 4°C with rocking. Beads were washed with lysis buffer and eluted with an imidazole step gradient (imidazole at 20, 40, 60, 80, 100, 250 mM) in elution buffer (20 mM NaPO_4_ pH 7.4, 300 mM NaCl, 5 mM β-mercaptoethanol). Protein used for biochemical experiments was dialyzed in FOG-3 buffer (20 mM HEPES pH 7.0, 50 mM NaCl, 0.5 mM (tris(2-carboxyethyl)phosphine) (TCEP)), while protein for crystallization was dialyzed in crystallization buffer (20 mM HEPES pH 7.0, 100 mM MgSO_4_, 0.5 mM TCEP). Samples were concentrated with Amicon Ultra-4 3000 MW concentrators (EMD Millipore) and run on a Superdex 200 (GE Healthcare). Recombinant protein was again concentrated with Amicon Ultra-4 3000 MW concentrators and protein concentration estimated by A280.

#### FOG-3 protease cleavage

Recombinant, full length FOG-3 with a C-terminal histidine tag was incubated with trypsin (Sigma-Aldrich) and elastase (Sigma-Aldrich) at room temperature prior to SDS-PAGE. The gel was stained with Coomassie to visualize cleavage products. Sample was also cleaved with trypsin or elastase for 45 minutes at room temperature (~20°C) and submitted for mass spectrometry (University of Wisconsin-Madison Biotechnology Center). The mass spectrometry fragment that most closely matched the SDS-PAGE band mapped to residues 1-135 for trypsin and 1-142 for elastase. Sequence alignments showed conservation up until residue 137, and thus we focused our structural efforts on this fragment.

#### Protein folding assay

The protein folding assay used followed published protocols^3^. Briefly, recombinant FOG-3 (1-137) with histidine tag was incubated with 90x concentrated SYPRO orange (5000x stock, Invitrogen) in FOG-3 buffer. 18 μl of the protein-dye mix was mixed with 2 μl Additive Screen (Hampton Research) and heated in a 7500 Real-Time PCR System thermocycler (Applied Biosystems) at 0.1°C/s from 20°C to 70°C while monitoring A405. SYPRO orange dye bound to unfolded protein. Thus, FOG-3 unfolded at a certain temperature, allowing dye binding and increasing A405 absorbance. The additive was judged as enhancing thermostability based upon the shift in the melting curve to the right, or requiring higher temperatures for signal. This assay was performed twice with similar results.

#### Crystallization, data collection, structure determination, and refinement

Crystallization conditions were screened with sitting drop trays set up by the Mosquito (TTP Labtech). We obtained crystals using recombinant FOG-3 (1-137 H48N, C117A) with an intact histidine tag and incubating our trays at 4°C. After 3 weeks, rhomboid crystals were observed in conditions A (0.1 M sodium citrate ph 5.6, 10% (vol/vol) isopropanol, 10% (wt/vol) PEG 4000) and B (0.1 M magnesium acetate, 0.1 M sodium citrate pH 5.6, 8% (wt/vol) PEG 10000). UV scanning with a UVEX-M (280 nm excitation, 350 nm emission; JANSi) identified these to be protein crystals. Both conditions were reproducible, but we were able to collect complete datasets from the crystals grown directly from the condition B screening trays. Phasing was accomplished with molecular replacement using Phaser^4^ and a human Tob homolog (PDB ID: 2Z15) as a starting model. Model building and refinement were done in Phenix^5^ and Coot^6^. Water molecules were first modeled by Phenix before checked manually. Three densities were too big to be water molecules. We could model one of the densities with sulfate. Two densities were observed in the solvent-accessible area adjacent to residues 52-56 in both copies in the ASU. Density is observed at FoFc contour levels past 6 sigma. We attempted modeling of acetate (too small) and citrate (too large), both present in the crystallization conditions, but the fit was unsatisfactory. Thus, the final uploaded model does not account for these two large densities. Coordinates and reflection data are available at RCSB (PDB ID: 5TD6).

#### Negative stain electron microscopy

Samples were negative stained with Nano-W (Nanoprobes, Yaphank, NY) using the two-step method. A 2 μl droplet of samples was placed on a Pioloform (Ted Pella) coated 300 mesh Cu Thin-Bar grid (EMS, Hatfield, PA), coating side down. The excess was wicked with filter paper and allowed to barely dry. A 2 μl droplet of Nano-W applied, wicked again with clean new filter paper, and allowed to dry. The sample was viewed on a Philips CM120 transmission electron microscope at 80 kV and documented with a SIS (Olympus / Soft Imaging Systems) MegaView III digital camera.

### Molecular Genetics

#### Worm maintenance

*C. elegans* were maintained as described previously^7^. Strains used: N2 Bristol

JK2739: *hT2[qIs48](I;III)/lin-6(e1466)dpy-5(e61)I*

JK4871: *fog-3(q520) I; qSi41[fog-3::3xFLAG] II*

JK5437: *fog-3(q847) I/hT2[qIs48](I;III)*

JK5439: *fog-3(q849) I/hT2[qIs48](I;III)*

Strains are available at the Caenorhabditis Genetics Center (cbs.umn.edu/cgc/home) or upon request.

#### *C. elegans* CRISPR

CRISPR of *fog-3* was achieved using a *dpy-10* roller co-injection strategy^8^. Briefly, an sgRNA construct containing the U6 promoter and sgRNA scaffold from pDD162^9^ along with the targeting sequence caatcagtccccgagtacg (pJK1910) and ggttctgaccacgtactcg (pJK1925) were cloned into the *Xma*I site of pUC19 using one step isothermal DNA assembly. The repair template was a 99 nt ssDNA oligo (ataaaaatactttaaatttcatttttccagctaccaatcagtccccAagtaTgtTgtcCgaaccgctgcaatccgcgcggagccttgctcgaatcttgg, IDT) that inserted an E126K mutation and removed an *Ava*I restriction site. Injections were performed in young N2 hermaphrodite *C. elegans,* using *fog-3* sgRNA plasmids, *dpy-10* sgRNA plasmid, *fog-3* E126K repair template, and Cas9 plasmid as described^8^, and F1 rollers were screened for the desired mutation by PCR and *Ava*I digest. Two alleles, *q847* and *q849*, were recovered from separate injected animals and therefore represent independent editing events. We verified the *fog-3* mutations by Sanger sequencing. Homozygous mutants had a Fog phenotype and thus could only produce oocytes. These worms were outcrossed twice with N2 before crossing with JK2739 containing balancer *hT2[qIs48](I;III)*.

#### Fertility and Fog phenotype

Heterozygous and homozygous *fog-3(q847),* or *fog-3(q849),* were singled onto plates as L4 larvae. After 3 and 4 days, worms were scored for the presence of L1 larvae and Fog phenotype. Four *q847* homozygous worms ruptured and died by day 3, and thus could not be properly scored.

#### iCLIP

*In vivo* crosslinking and immunoprecipitation (iCLIP) was carried out essentially as described^10^, with modifications to worm growth, crosslinking, lysis and RNase digestion described here.

#### Nematode culture and UV crosslinking for iCLIP

*C. elegans* strain JK4871 L1 larvae were obtained by bleaching and synchronizing by standard methods^11^. Larvae were plated onto 10 cm OP50 plates (~50,000 per plate) and propagated at 20°C for ~40-46 hours until most of the worms were at the early L4 stage when FOG-3 expression is greatest. Worms were washed with M9 (42.3 mM Na_2_HPO_4_, 22 mM KH_2_PO_4_, 85.6 mM NaCl, 1 mM MgSO_4_), pooled into 250,000 worm samples, and placed on a 10 cm NGM agarose plate. Liquid was removed from the plate as much as possible. Animals were irradiated two times sequentially at 254 nm with 0.9999 J/cm^2^ in a XL-1000 UV Crosslinker (Spectrolinker). Non-crosslinked samples were incubated at room temperature as a negative control for the radiolabeled gel (Fig 3). For the iCLIP negative control, we performed the pulldown of crosslinked JK4871 worm lysate with beads alone (no antibody). Worms were rinsed from the plates with cold M9, washed once, and transferred to a 2 mL Eppendorf tube. The pellet was washed again in freezing buffer (50 mM Tris pH 7.5, 150 mM NaCl, 10% (wt/vol) glycerol, 0.05% (vol/vol) tween 20) and frozen with liquid nitrogen. Pellets were stored at -80°C until use.

#### Lysis and RNA digestion

*C. elegans* pellets were thawed by adding ice cold lysis buffer (50 mM Tris pH 7.5, 100 mM NaCl, 1% Pierce NP-40, 0.1% SDS, 0.5% sodium deoxycholate, Roche cOmplete EDTA-free Protease Inhibitor Cocktail, Ambion ANTI-RNase) and incubated for 20 minutes at 4°C with rocking. The thawed pellets were centrifuged at 1,000 x *g*, 4 °C for 1 minute and washed 3 times with ice cold lysis buffer. Lysis buffer was added to the pellet along with a 5 mm stainless steel ball (Retsch). Lysis was performed in the cold room using a 400 MM mill mixer (Retsch). Lysis was completed after three 10 minute cycles at a setting of 30 Hz, with four-minute freeze-thaws after the first and second cycles. Freeze-thaws were performed by immersion in liquid nitrogen for 1 minute, then returning to liquid state by immersion in room temperature water for 4 minutes. Worm lysis was confirmed by observing a small aliquot of final lysate on a dissection scope. The lysate was cleared by centrifugation for 15 minutes at 16,100 x *g*, 4°C. Protein concentration of the cleared lysate was determined with the Direct Detect spectrometer (EMD Millipore). Our pellets containing 250,000 worms yielded ~12 mg/mL of total protein, and we used 10 mg total protein per biological replicate. Double RNase digestion of protein-RNA complexes was performed as previously described^12^. For the first digestion, which occurred immediately after lysis and just prior to immunoprecipitation, guanosine specific RNase T1 (ThermoFisher) was added to the cleared lysate at a final concentration of 1 Unit/μL. The sample was incubated in a Thermomixer for 15 minutes at 22 °C, 1100 rpm and then cooled on ice for 5 minutes.

Protein G Dynabeads (Life Technologies) were aliquoted to a fresh RNase-free roundbottom tube (USA Scientific). The tube was placed on a Dynal magnet (Invitrogen), the existing buffer was removed, and M2 FLAG antibody (Sigma-Aldrich) added at 20 μg antibody to 3 mg Protein G dynabeads in PBS-T (PBS pH 7.2 (137 mM NaCl, 2.7 mM KCl, 10 mM Na_2_HPO4, 1.8 mM KH_2_PO4), 0.02% Tween-20). The beads plus antibody solution was incubated at room temperature on a rotator for 45 minutes. The tube was again placed on the magnet, the antibody solution removed, and the cleared lysate was added. Immunoprecipitation was carried out overnight at 4 °C. As a negative control, we performed the pulldown of crosslinked JK4871 worm lysate with beads alone (no antibody).

Following immunoprecipitation, the beads were washed as described^10^, with minor modifications. We performed washes in the cold room (~4 °C) with two wash buffers: a high-salt wash buffer (50 mM Tris-HCl pH 7.5, 1 M NaCl, 1 mM EDTA, 1% Pierce NP-40, 0.1% SDS, 0.5% sodium deoxycholate) and PNK buffer (20 mM Tris-HCl pH 7.5, 10 mM MgCl2, 0.2% Tween-20). The second RNase T1 digestion was then performed on the washed beads at a final concentration of 100 Units/μL in PNK buffer. Samples were incubated in a Thermomixer for 15 minutes at 22 °C shaking at 1100 rpm, cooled on ice for 5 minutes, and then processed through the remaining iCLIP protocol as described^10^. We confirmed immunoprecipitation of FOG-3 from experimental versus negative control samples by immunoblot with an M2 FLAG antibody (**Supplementary Fig. 4b**). We confirmed that 3xFLAG-FOG-3 crosslinked to RNA by visualizing 5’ radioactively labeled RNA bound to the FOG-3 protein when antibody was present on the beads (**Fig. 3d** and **Supplementary Fig. 4d**).

Single-end sequencing was performed on an Illumina HiSeq 2000 (University of Wisconsin Biotechnology Center). The cDNA library of each replicate was prepared with a unique “Rclip” reverse transcription primer (as in Huppertz, et al.^10^), which contained a partially randomized sequence (i.e., a “barcode”). The constant portion of the barcode enabled each read to be identified by replicate and allowed for replicate multiplexing. The randomized portion of the barcode allowed for computational filtering of artifacts from individual reads caused by PCR amplification of the cDNAs, such as read duplication. After high-throughput sequencing, the barcode sequence preceded the cDNA sequence and thus could be easily identified and removed prior to read mapping.

#### iCLIP sequence analysis

Reads (**Supplementary Table 2**, Tab 1, column B) were aligned to the WS235 genome using STAR ^13^ and previously described parameters^14^, except for the parameter –alignEndsType Local (mismatches at the ends of reads are tolerated). Multimapping reads were removed, and high-confidence mappings were selected as those with alignment scores of at least 20 (**Supplementary Table 2**, Tab 1, column C). PCR duplicates were collapsed to unique reads (**Supplementary Table 2**, Tab 1, column E) using the method described in Weyn-Vanhentenryck, et al.^15^. Reads were assigned to genes using HTSeq^16^. CIMS (crosslinking induced mutation sites) and CITS (crosslink induced truncation sites) analyses were performed as described previously ^15^, except we did not require CIMS to reproduce between replicates, and are included in **Supplementary Table 3**, tab 3. For peak analysis, “clusters” were defined as regions of overlapping reads. Using the reads indicated in **Supplementary Table 2**, tab 1 (column E), all reads within a gene had their position randomized 1000 times to empirically determine a cluster p value as the odds of having a cluster with the given maximum read depth from randomized read positions. This is similar to the local Poisson method ^17^, as the Poisson approximates of read scrambling. A Benjamini-Hochberg (BH) correction for multiple hypothesis testing was then applied at 1% FDR, resulting in the cluster numbers in **Supplementary Table 2** (Tab 2, column F). Finally, only overlapping clusters called independently as significant in at least 2 of the 4 replicates were retained as reproducible clusters, resulting in the cluster numbers in **Supplementary Table 2** (Tab 2, column G). Final clusters for FOG-3 and control samples are given **Supplementary Table 3**. While this is a simple method that does not account for background RNA abundance, it resulted in only 6 clusters for the negative control samples, suggesting it is effective at removing background in our datasets. We define peaks as all maxima at least 5 reads deep and at least 5% of the highest peak in the given gene; we counted neighboring peaks as distinct only if signal dropped to 50% or less of the lower peak maxima. Our definition of peaks differs from our definition of clusters, which are regions of continuous read coverage that pass the 1% FDR threshold. Clusters extend until iCLIP coverage drops to zero, thereby containing any number of distinct signal concentrations, and motivating our separate definitions of peaks and clusters.

To compare our results with previous FOG-3 RIP results^18^, we calculated overlap with the top 722 FOG-3 targets, and evaluated significance by Fisher’s exact test. To determine whether FOG-3 targets were associated with oogenesis, spermatogenesis, or mitosis, we used the method described previously^18^, with significance evaluated by Fisher’s exact test. Figures depicting iCLIP results (**Supplementary Fig. 4i,j**) were generated using Matplotlib^19^, and python scripts available at https://github.com/dfporter. Raw sequence files of all replicates are available through Gene Expression Omnibus (GEO accession GSE76521).

---

**Supplementary Table 1.** Data collection and refinement statistics.

|  | ***C. elegans* FOG-3** |
| --- | --- |
| **PDB ID** | 5TD6 |
| **Wavelength (Å...)** | 0.9537 |
| **Resolution range (Å...)** | 29.22 - 2.034 (2.106 - 2.034) |
| **Space group** | P 31 2 1 |
| **Unit cell** | 64.992 64.992 133.57 90 90 120 |
| **Total reflections** | 147208 (14689) |
| **Unique reflections** | 21617 (2091) |
| **Multiplicity** | 6.8 (7.0) |
| **Completeness (%)** | 99.79 (98.18) |
| **Mean I/sigma(I)** | 24.12 (2.56) |
| **Wilson B-factor** | 39.54 |
| **R-merge** | 0.0498 (0.7383) |
| **R-meas** | 0.05399 |
| **CC1/2** | 1 (0.789) |
| **CC\*** | 1 (0.939) |
| **Reflections used for R-free** | 1993 |
| **R-work** | 0.1769 (0.2255) |
| **R-free** | 0.2285 (0.2886) |
| **Number of non-hydrogen atoms** | 2210 |
| **macromolecules** | 2067 |
| **ligands** | 5 |
| **water** | 138 |
| **Protein residues** | 260 |
| **RMS(bonds)** | 0.009 |
| **RMS(angles)** | 1.10 |
| **Ramachandran favored (%)** | 98 |
| **Ramachandran allowed (%)** | 2 |
| **Ramachandran outliers (%)** | 0 |
| **Clashscore** | 3.89 |
| **Average B-factor** | 46.60 |
| **macromolecules** | 46.70 |
| **ligands** | 32.50 |
| **solvent** | 45.50 |

Statistics for the highest-resolution shell are shown in parentheses.

